# Prediction of Optimal Growth Temperature using only Genome Derived Features

**DOI:** 10.1101/273094

**Authors:** David B. Sauer, Da-Neng Wang

## Abstract

Optimal growth temperature is a fundamental characteristic of all living organisms. Knowledge of this temperature is central to the study the organism, the thermal stability and temperature dependent activity of its genes, and the bioprospecting of its genome for thermally adapted proteins. While high throughput sequencing methods have dramatically increased the availability of genomic information, the growth temperatures of the source organisms are often unknown. This limits the study and technological application of these species and their genomes. Here, we present a novel method for the prediction of growth temperatures of prokaryotes using only genomic sequences. By applying the reverse ecology principle that an organism’s genome includes identifiable adaptations to its native environment, we can predict a species’ optimal growth temperature with an accuracy of 4.69 °C root-mean-square error and a correlation coefficient of 0.908. The accuracy can be further improved for specific taxonomic clades or by excluding psychrophiles. This method provides a valuable tool for the rapid calculation of organism growth temperature when only the genome sequence is known.

## Author Summary

The optimal growth temperature is a fundamental characteristic of all living organisms. It is the temperature at which the organism grows at the greatest rate, and is a consequence of adaptations of that organism to its native environment. These adaptations are contained within the genome of the organism, and therefore species from varying environments have distinct genomic characteristics. Here we use those genomic characteristics to predict a species’ optimal growth temperature. This provides a novel tool for describing a key parameter of the species’ native environment when it is otherwise unknown. This is particularly valuable as the rate of genome sequencing has increased, while the determination of growth temperature remains laborious.

## Introduction

Growth conditions of an organism are essential to its characterization. However, these values may be unknown in organisms which are difficult to culture, “unculturable”, or otherwise poorly characterized. Reverse ecology posits that the evolutionary effects of an organism’s native environment is reflected by adaptations in its genome [1]. Therefore, an organism’s native environment can be identified by comparing its genome to the genomes of other organisms from a range of environments. Notably, this is done without experimental manipulation or interrogation of the organism beyond genome sequencing. Such reverse ecology strategies have been successful in studying adaptation to soil conditions [2], salinity [3], and temperature [4].

Of these environmental pressures, temperature, being a description of the internal energy of the environment, is a particularly strong driving force for adaptation. Prokaryotes are often viable over a range of temperatures, which varies by species. For a particular organism, increasing temperature beyond it’s growth range, corresponding to increased internal energy, can lead to loss of protein and nucleic acid structure. Conversely, a sub-optimal temperature leads to reduced enzyme kinetics and stiffening lipid membranes. Each of these biological consequences may be deleterious to un-adapted organisms. Therefore, it is perhaps not surprising that an organism’s optimal growth temperature (OGT) correlates to quantifiable properties (features) in the organism’s nucleotide and protein sequences. Features correlated with OGT can be identified in the genomic [5], tRNA [6,7], rRNA [6–8], open reading frame [9,10], and in the proteomic sequences [10–13]. Correlations between OGT and tRNA G+C content [6,7] or the charged versus polar amino acid ratios [14] are well known.

Clearly, OGT is a necessary parameter for analyzing physiological processes of an organism or activities of its genes and proteins. [15,16]. However, the experimental determination of OGT is laborious [17,18], and sometimes unattainable [19]. Also, recorded OGT or environmental temperature may be inconsistently measured, particularly in genetic samples not obtained from pure culture [20]. Further, for metagenomic samples the conditions during collection may significantly differ from the originating species’ growth environment. This can be due to the organism or its genetic material being found distant from its originating environment [21], or the collected genomic material may be from organisms which are inviable [22]. Even in pure culture in the laboratory, the experimental growth conditions can vary greatly [23] and may not be at the source organisms’ OGT [24].

While many previous studies have aimed to identify genes and proteins [25], mutations [16], and mechanisms [15] that drive thermal adaptation, there is also great value in using these adaptive differences to provide data of an organism’s native environment when it may not be otherwise known or well-described. A number of parameters have been identified which correlate with OGT [14]. However, those correlations are often weak and therefore of limited predictive value alone. Here, we aim to predict a prokaryotic species’ OGT only from its genomic sequence. We set out to develop a novel tool for the ecological characterization of a species based solely on its genome, the study of thermoadaptation, and bioprospecting for thermoadapted genes.

## Results

### Prokaryote genome redundancy is highly skewed

Of the initial 8270 prokaryotic species with a reported OGT, genome sequences were available for 2708 species. These sequenced species were composed of 2538 Bacteria and 170 Archaea, with OGTs ranging from 4 to 103 °C. A total of 36,721 sequenced genomes for these species were downloaded, indicating multiple genomes for each species on average. However, the number of genomes per species was highly skewed, with great redundancy for model organisms and pathogens (Fig S1C). To avoid having these relatively few species dominate the analysis, features were averaged by species and all regressions were done by species rather than by genome.

### Individual genome derived features correlate with OGT

Based on the reverse genomics principle that an organism’s adaptation to its environment is reflected within its genome, we hypothesize that a species’ OGT could be predicted based on characteristics of its genome and genome derived sequences. This hypothesis was supported by previous noted correlations between OGT and individual features of the genomic [5,6,26], tRNA [6,7], rRNA [6–8], open reading frame [7,9,10,27,28], and proteomic or protein sequences [10–14,29–35]. These features are quantifiable properties of the sequence, such as G+C content, length, and nucleotide or amino acid fraction. Of the features calculated, 42 were found in this work to be correlated with OGT in the present dataset by the Pearson correlation coefficient with |r| > 0.3 (Fig 1, Table S1). However, these correlations to OGT were often weak and therefore insufficient for the calculation of a species’ growth temperature. Furthermore, there was a strong association among many features (Fig S2). We therefore decided to consider them simultaneously, using multiple linear regression, with features added individually to minimize multicollinearity. We started by classifying features based on the source sequences (genomic, tRNA, rRNA, open reading frames, and proteome). Multiple linear regressions were calculated, progressively increasing the number of feature classes used in the regression.

**Figure 1.**
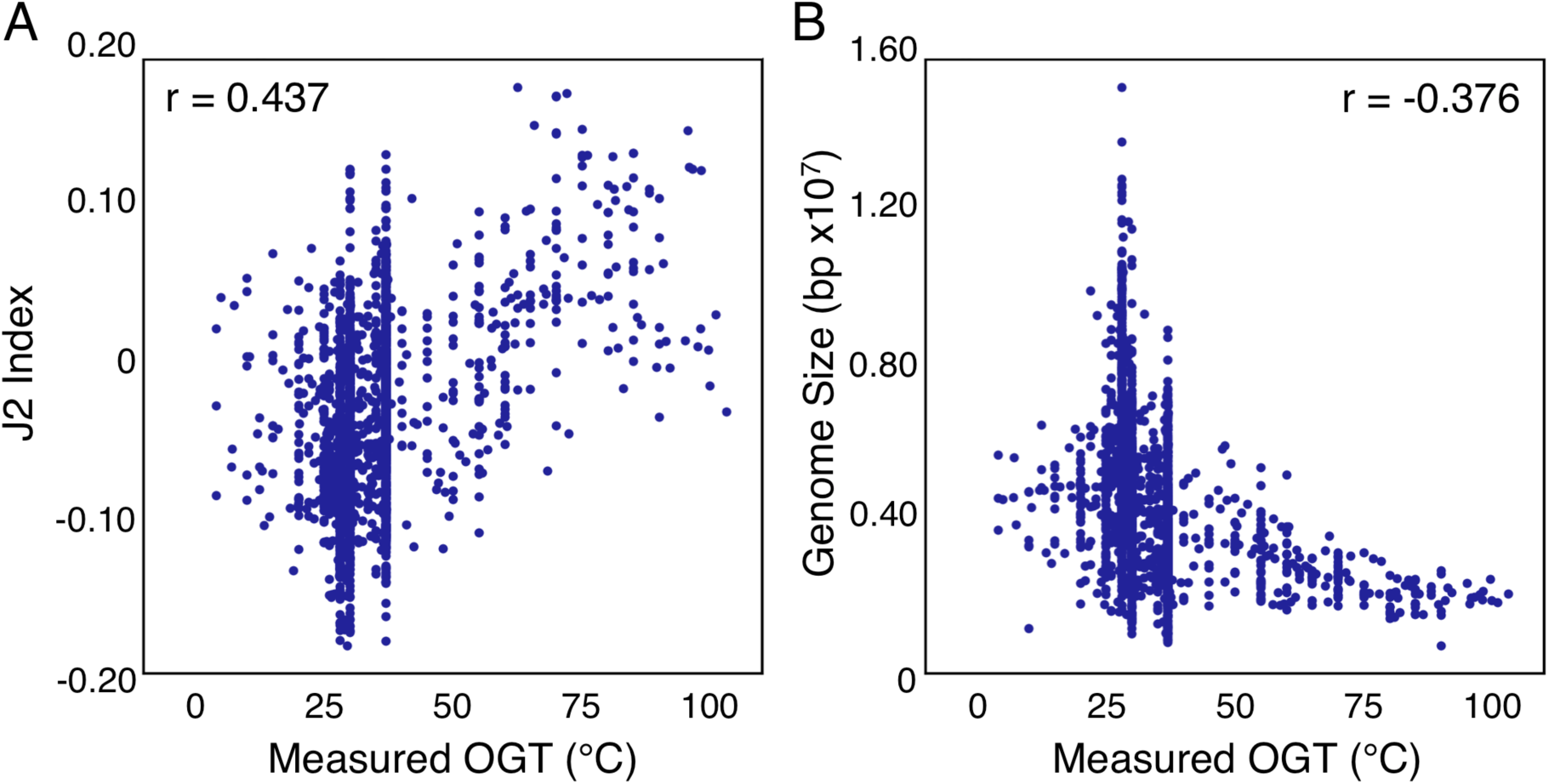
Individual genome derived features correlate weakly with the originating species’ OGT. Measure optimal growth temperature for each species versus J2 index of genomic dinucleotide fractions (A) and total genome size (B).

### A regression using only genomic sequence based features is weakly predictive of OGT

The genomic sequence provides information about the nucleotide content, nucleotide order, and chromosomal structure of an organism’s hereditable genetic material. In the absence of any other knowledge, this sequence still reflects adaptations to the particular thermal environment of the organism. For example, total genome size has been shown to be negatively correlated with a species’ OGT [26]. Accordingly, it has been proposed that the reduced time and energy of genomic replication offers selective advantages at higher temperatures. Additionally, the necessity of maintaining genomic structure with increased temperature is thought to be reflected in a species’ genomic dinucleotide fractions [36], which is quantified in the J2 index [5]. In the present dataset, individual nucleotide and dinucleotide fractions of the genome, the J2 index, the G+C content, and total size were calculated for each genome. Of these features, the J2 index, genome size, and the CT and AG dinucleotide fractions correlated with OGT, but only weakly. Using these poorly correlated and collinear input features for regression, the resulting multiple linear regression is poor at predicting OGT with a root mean squared error (RMSE) of 9.86 °C (r = 0.469) (Fig S3).

### tRNA and rRNA sequences improve OGT prediction

tRNA and rRNA are nucleic acids whose structure, and enzymatic activity in the case of rRNA, are essential to cell viability. Therefore, the direct correlation of OGT to G+C content of tRNAs [6,7] and rRNAs [8,37] is thought to reflect the necessary increase in base pair hydrogen bonding needed to maintain the structure of these nucleic acids at elevated temperatures. While a subset of the previously analyzed genomic sequence, we hypothesized that features derived from these tRNA and rRNA sequences might be more strongly correlated with OGT. To this end, we identified their sequences bioinformatically. tRNA and 16S rRNA sequences were identified in 100% and 98% of the species respectively, reflecting the highly conserved nature of these genes.

Using these identified tRNA and rRNA sequences, nucleotide fractions and G+C content were calculated for each. All calculated features for tRNA and rRNA sequences were correlated with OGT. Calculating a new linear regression with the OGT using tRNA features, in addition to genomic features, improved accuracy (RMSE = 7.30 °C, r = 0.757) (Fig 2A). Similarly, a regression calculated with rRNA and genomic features also improved accuracy (RMSE = 6.99 °C, r = 0.784) (Fig 2B). By using all available tRNA, rRNA, and genomic features, a still more accurate linear regression was calculated (RMSE = 6.71 °C, r = 0.802) (Fig 2C).

**Figure 2.**
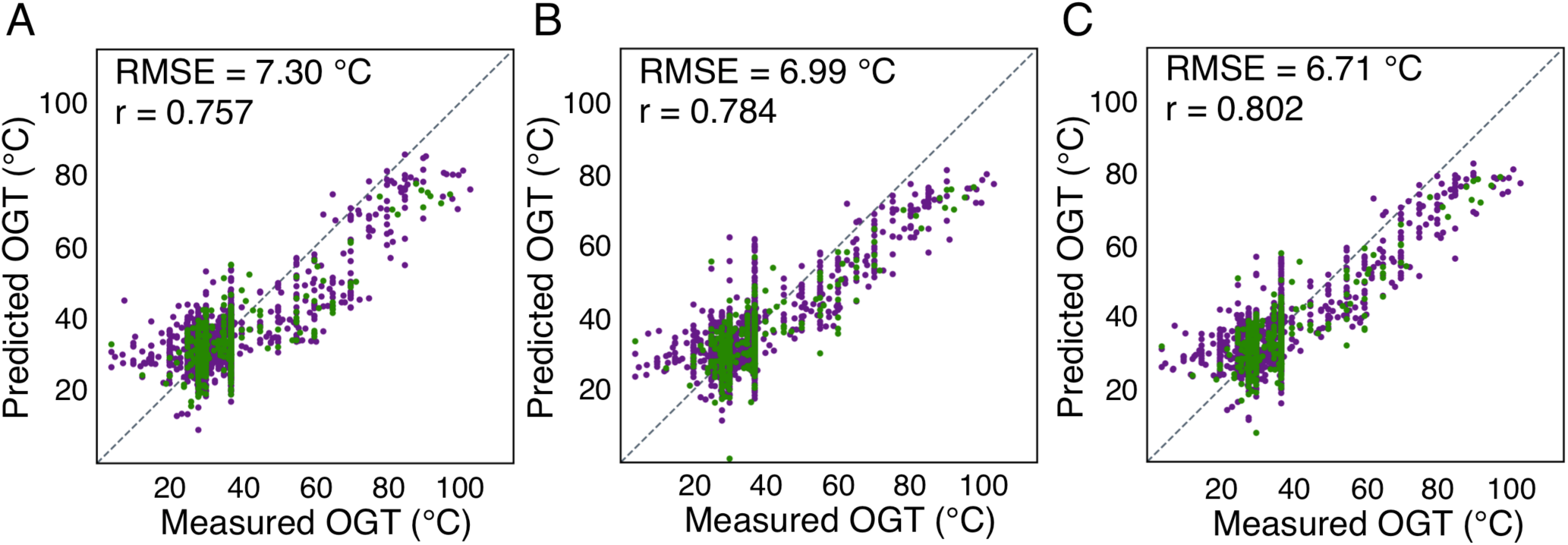
Using genomic and genic sequences improve OGT prediction accuracy. Predicted versus measured OGT for each species, using linear regressions with features derived from genomic and (A) tRNA, (B) rRNA, or (C) tRNA and rRNA sequences. Species used for regression and evaluation are shown in purple and green, respectively. The dotted line indicates a perfect prediction.

### ORF sequences improve OGT prediction

As tRNA and rRNA features clearly improve the ability to predict a species’ OGT, we examined if other gene sequences might also improve the regression. In particular open reading frames, which code for proteins but exclude the non-coding regions of the genome, were considered. We hypothesized that using coding regions alone would increase sensitivity to changes in OGT. Additionally, codon biases have previously been reported to correlate with OGT [13], likely reflecting both amino acid differences and the necessity of maintaining proper codon-anticodon pairing in differing thermal environments. Furthermore, the greater number of ORFs in a genome, relative to tRNAs and rRNAs, make the features of ORFs less sensitive to single gene aberrations or mispredictions. Therefore, ORF derived features were hypothesized to more sensitively and accurately report on the thermal environment than tRNA or rRNA sequences.

We identified ORFs within the genomic sequences bioinformatically. From these ORFs, a number of derived features were calculated including nucleotide and dinucleotide fractions, codon fractions, start and stop codon fractions, the coding ratio and fraction of the genome, the ORF density of the genome, G+C and A+G content, and average length. Of these, nine were found to be correlated with OGT. These include the A+G content, codon and dinucleotide fractions, and the fraction of the alternative start codon TTG. These ORF derived features, in addition to the genomic, tRNA, and rRNA features, were used to calculate a new multiple linear regression with significantly improved accuracy (RMSE = 5.77 °C, r= 0.857) (Fig 3).

**Figure 3.**
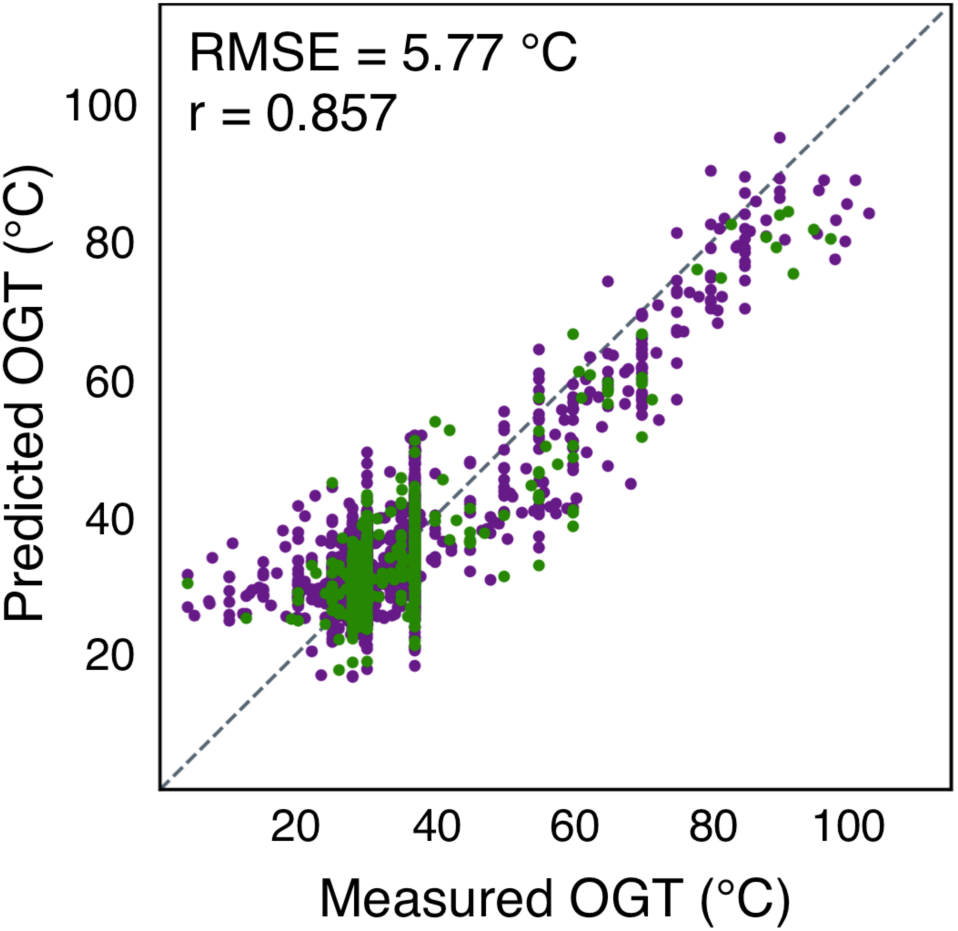
Open reading frame sequences further improve OGT prediction accuracy. Predicted versus measured OGT for each species, using a linear regression with features derived from sgenomic, tRNA, rRNA, and ORF sequences. Species used for regression and evaluation are shown in purple and green, respectively. The dotted line indicates a perfect prediction.

### Including proteome features significantly improves OGT prediction

While ORF feature correlation to OGT partially reflects the adaptation of the coding regions and mRNAs to the thermodynamic environment, it has been suspected that this correlation also reflected adaptations in each species’ proteome to OGT. Temperature is known to correlate with protein folding, biochemistry, and enzyme kinetics, all of which are essential to organismal viability [10,14,32]. Based on these biological consequences, proteome derived features were hypothesized to be especially sensitive to thermal environment. Therefore, the proteome was translated from each species’ ORFs, and features calculated from the proteome’s primary sequence. These features included amino acid fractions, the fraction of the proteome to be charged or thermolablile, and the EK/QH, LK/Q, Polar/Charged, and Polar/Hydrophobic amino acid ratios.

Supporting this hypothesis, proteome derived features were found to have the strongest correlation to OGT (Table S1), with the greatest correlation being the fraction of the proteome composed of the amino acids ILVWYGERKP [13]. The linear regression of OGT using proteome features, in addition to previously described features, significantly improved accuracy (RMSE = 4.69 °C, r = 0.908). (Fig 4, Eq S1).

**Figure 4.**
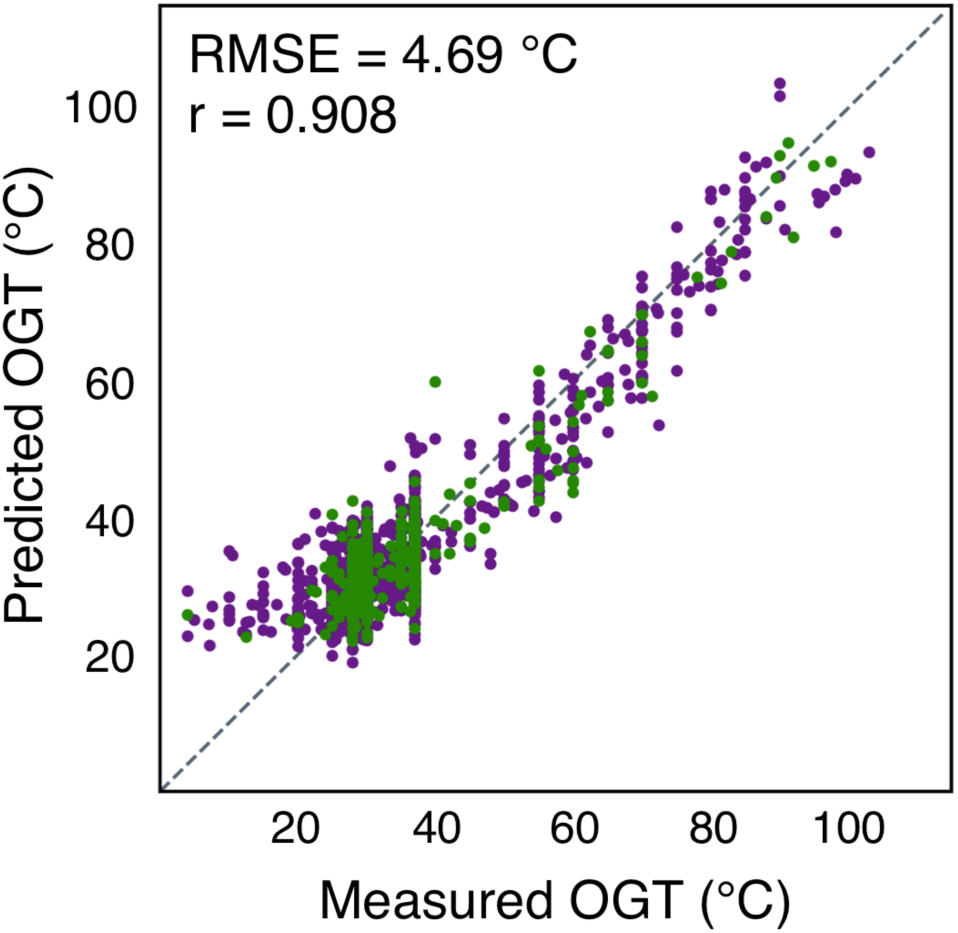
Proteome derived features significantly improve OGT prediction accuracy. Predicted versus measured OGTs for each species, using a linear regression with features derived from genomic, tRNA, rRNA, ORF, and proteome sequences. Species used for regression and evaluation are shown in purple and green, respectively. The dotted line indicates a perfect prediction.

### Taxonomic clade specific regressions are the most accurate

The regressions described up to this point were all made using all prokaryotic species. However, we had noted that the number of individual features correlated with OGT was much higher in Archaea than Bacteria (Table S1). In addition, we hypothesized that the magnitude of the response of each feature to OGT may be distinct in each superkingdom.

Based on these distinctions, we tested whether superkingdom specific regressions would be more accurate than the regression of all prokaryotes (Fig. 5). Using the NCBI taxonomic assignment for each species, an Archaea-only regression dramatically improved accuracy for these species (RMSE = 3.21 °C, r = 0.995) (Eq S2). However, the Bacteria-only regression only showed only a slight improvement (RMSE = 4.61 °C, r = 0.816) (Eq S3). This likely reflects bias of the general prokaryotic regression, due to the numerical majority of bacterial species and the greater diversity of bacterial species.

**Figure 5.**
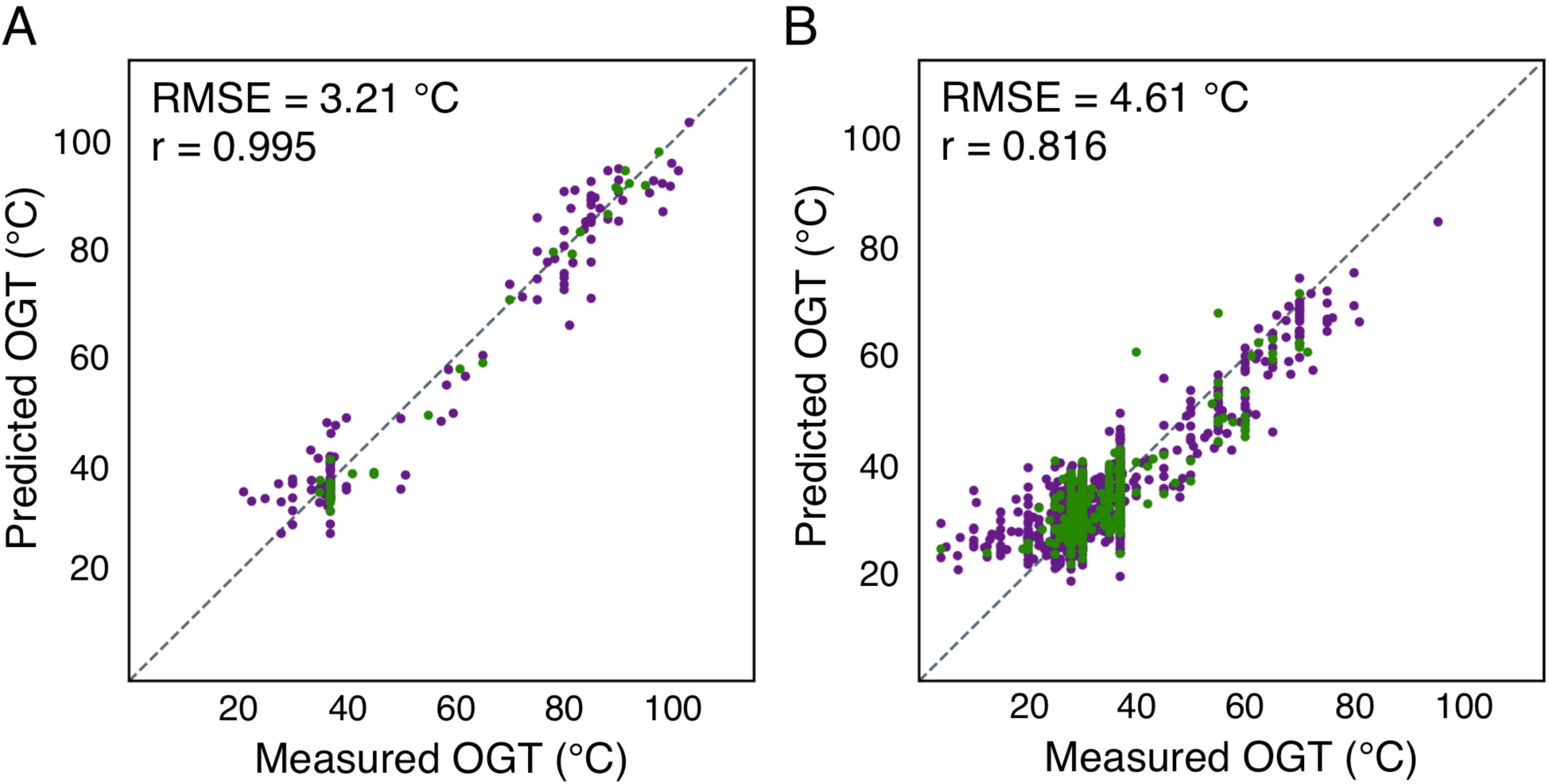
Taxon specific linear regressions are most accurate. Predicted versus measured OGT for each species using superkingdom specific linear regressions for Archaea (A) and Bacteria (B). Species used for regression and evaluation are shown in purple and green, respectively. The dotted line indicates a perfect prediction.

Addressing this diversity in bacteria, the taxonomic specific regression can be further improved when the data is separated by phylum or class. OGT regression was limited to clades where the number of species (N) was greater than 50 to ensure the significance of the regression. Of the individual phyla, the most accurate regressions are found in the *Firmicutes* (RMSE = 4.88 °C, r = 0.831), *Actinobacteria* (RMSE = 2.90 °C, r = 0.818), *Bacteroidetes* (RMSE = 1.58 °C, r = 0.964), and *Euryarchaeota* (RMSE = 4.00 °C, r = 0.985) (Fig S4). In contrast, the *Proteobacteria* regression had much more weakly correlated predicted and reported OGTs (RMSE = 4.10 °C, r = 0.569), though the small RMSE likely reflects the narrow OGT range of this phylum. Further subdivision of the *Proteobacteria* into classes (Fig S5) resulted in significant correlation of the *Betaproteobacteria* (RMSE = 2.94 °C, r = 0.789), and *Deltaproteobacteria* regressions (RMSE = 2.04 °C, r = 0.761). However, no correlation was found in regressions for the *Proteobacteria* classes of *Alphaproteobacteria* or *Gammaproteobacteria*.

### Discussion

Knowing an organism’s optimal growth characteristics is central to addressing basic biological questions about how organisms adapt to a particular environmental niche. Further, the systematic study of adaption often requires the optimal growth conditions of the species of origin for each species and gene or protein examined. Additionally, proteins from organisms adapted to particular environmental niches are often particularly suited for structural biology [38–40] and industrial applications [41,42].

However, if the growth characteristics of already sequenced organisms are uncharacterized, the physiochemical properties of these genes that otherwise might be inferred are lost [20]. Consequently, this limits the use of these genomes in academic study and mining for biotechnology applications. Exacerbating this issue, high throughput sequencing has enabled rapid growth in the number of available genomic, metagenomic, and derived proteomic sequences. This growth in genetic information is likely to outpace the laborious experimental task of characterizing the growth conditions of each species, leading to an increasing number of genomic sequences with unknown growth characteristics. This is already apparent by those organisms which have been ‘unculturable’ to date, but which have been sequenced by metagenomics.

To satisfy the need for growth condition data when only genomic sequences are available, here we demonstrate a novel reverse ecology tool to accurately predict the OGT using solely the genomic sequence as input. Our method can predict the OGT for sequenced Archaea and bacteria with an accuracy of 3.21 °C and 4.61 °C, respectively.

### OGT can be accurately predicted using only genome derived parameters

Genome classification is clearly essential to the most accurate prediction of OGT. The programs used for tRNA, rRNA, and ORF identification all require some level of taxonomic classification. When applying the general prokaryotic regression, this is only requires the relatively simple exclusion of eukaryotic samples prior to sequencing [43]. However, the most accurate OGT regressions are taxon specific, and therefore genomic samples require further classification. This assignment is routinely addressed *in silico*, using specialized bioinformatic tools which can easily assign taxonomic clade to genomic material [44,45].

As a simple proof-of-concept, the prokaryotic genomes were also classified by superkingdom using the best scoring 16S rRNA hidden Markov model in Barrnap (Fig S6). These regressions were of similar accuracy to those using NCBI superkingdom assignments.

### Excluding genome size does not alter the regression accuracy

While prokaryote genome size is strongly correlated with OGT, it is unique among all features used here in requiring a complete genome for calculation. Therefore, this feature might not be available in metagenomic samples, or otherwise incompletely assembled genomes. Excluding this feature has only a minor impact on the regression for all prokaryotes (RMSE = 5.07 °C, r = 0.891), or the separate regressions for Bacteria (RMSE = 4.97 °C, r = 0.783), or Archaea (RMSE = 3.21 °C, r = 0.995) (Fig S7).

### Psychrophiles are poorly fit

While the final regressions of prokaryotes and Bacteria were generally accurate, species with optimal growth temperatures less than approximately 25 °C are clearly poorly fit. This outcome is unsurprising, as few psychrophilic sequences are present in the dataset (Fig. S1), and the mechanisms of thermoadaptation to higher and lower temperatures are not equivalent [46]. Excluding those species with an OGT of less than 25 °C yields a slightly better general prokaryotic regression (RMSE = 4.42 °C, r = 0.916) (Fig. S6). The archaeal regression was slightly worse (RMSE = 3.12 °C, r = 0.993), while the bacterial regression improved (RMSE = 4.26 °C, r = 0.832), reflecting the known OGT ranges of each superkingdom.

### Improvements over comparable methods

Our method significantly expands and improves upon the individual features previously described to correlate with OGT. By studying a much larger set of genomes, a more precise correlation between each feature and OGT can be calculated. Further, by using multiple features, more accurate and predictive regression models have been calculated. Notably, our method improves on previously reported analyses requiring particular genes being present in the genome, thereby making the method more general in application [47]. Also, this method quantitatively predicts an OGT rather than using classification (psychrophile, mesophile, thermophile, or hyperthermophile). This improves on methods which predict OGT ranges [47–50], where classification necessarily limited accuracy.

The most comparable method is reported by Zeldovich *et al.* calculating OGT from the proteome as OGT = 937F – 335, where F is the sum of the proteome fraction for the amino acids IVYWREL [13]. Using the current larger dataset, we calculate a lower correlation (r = 0.726) and accuracy (RMSE = 10.5 °C) than previously reported. This is likely a consequence of more genomic sequences being available, and our keeping of individual species separate rather than averaging those with the same OGT. By considering more features derived from the source organism’s genome, the prokaryotic regression presented here clearly advances upon this previous method improving in both correlation and accuracy. While we focus on growth temperature, the same principle could be readily applied to other quantifiable characteristics of an organism’s optimal growth environment, such as pH, salinity, osmolarity, or oxygen concentration.

### Application and validation

Applying these regressions, we predicted OGTs for those species with a genomic sequence available, but without a reported OGT in Sauer *et al.* (2015), using the most taxon specific linear regression available. Only the *Betaproteobacteria* and *Deltaproteobacteria* classes of *Proteobacteria* were predicted, excluding the *Alphaproteobacteria*, *Gammaproteobacteria*, and other *Proteobacteria* due to the poor predictive values of those taxon specific regressions. In total, 482 species’ OGTs were predicted (Table S2). Of the species with newly predicted OGTs, a more recent literature search revealed reported OGTs for 36 species [51–87]. The predicted and measured OGTs were strongly correlated (RMSE = 6.94 °C, r = 0.857), validating the predictive value of this method (Fig S9).

## Materials and Methods

### Source data and sequence extraction

Experimentally measured OGTs of various prokaryotic species were used as previously published without modification [88]. Taxonomic assignments for each species were collected from NCBI [89]. All available top level genome sequences for each species were downloaded from Ensembl [90]. tRNA sequences were identified with tRNAScan-SE 1.3.1 [91] with general settings. Ribosomal RNA genes were identified with Barrnap 0.8 [92] using superkingdom specific hidden Markov models, and rRNA sequences extracted from the genome using BEDtools 2.26.0 [93]. Open reading frame sequences were identified with GenemarkS 4.32 [94] using the default settings. ORFs were also translated into protein sequences using the standard genetic code. Features were calculated for each genome and derived proteome, ignoring ambiguous nucleotides and amino acids. All calculated features were averaged by species. Twenty percent of the species with available genomes were set aside as a test set and never used for regression, only evaluation.

### Multiple linear regression

Only individual features linearly correlated with OGT (|r| > 0.3) were used for multiple linear regression. To minimize multicollinearity, the initial regression input feature set consisted of only the feature most correlated with OGT. To this set all other correlated features were added individually, and multiple linear regressions were calculated. If the correlation between measured and predicted OGTs increased for any regression, the input feature which most increased the correlation was added to the input set. This was repeated until the correlation did not increase.

### Regression evaluation and prediction

The test set was only used for evaluation of the multiple linear regressions, comparing the calculated and measured OGTs. Regressions were evaluated by comparing the predicted and reported OGT using the Pearson correlation coefficient and root mean square error.

### *De novo* OGT prediction and validation

All top level genomes in Ensembl Bacteria were downloaded for each species where there was not a reported OGT in the Sauer *et al.* (2015) dataset. Taxonomic assignment and feature calculation were preformed as described above. The most taxonomic specific regression available, using genomic, tRNA, ORF, and proteome features, was used to predict the OGT for each species. For these newly predicted species, Pubmed was searched using the binomial name and “optimal growth” as keywords. From the returned publications, OGTs were manually collected where available.

Analyses were carried out using custom Python scripts using Biopython 2.7.12 [95], NumPy 1.13.3 [96], SciPy 1.0.0, Scikit-learn 0.19.1 [97], and MatPlotLib 2.1.0 [98].

## Acknowledgements

The authors thank Jennifer Marden for discussion and critical review of this manuscript.

This work was financially supported in part by an American Cancer Society Postdoctoral Fellowship (16-A1-00-005739 to D.B.S), the Department of Defense (W81XWH-16-1-0153 to D.B.S.), and NIH (R01-GM121994, R01-DK099023, and R01-GM093825 to D-N.W). This work was supported by the Office of the Assistant Secretary of Defense for Health Affairs, through the Peer Reviewed Cancer Research Program under Award No. W81XH-16-1-0153. Opinions, interpretations, conclusions and recommendations are those of the author and are not necessarily endorsed by the Department of Defense.

## Supporting Information Captions

Figure S1. The genomes available are dominated by mesophiles, bacteria, and repetitively sequenced organisms.

Figure S2. Features are often highly associated.

Figure S3. Using only genomic sequence features is poorly predictive of OGT.

Figure S4. Phylum specific regressions are often strongly predictive.

Figure S5. Class specific regressions can be strongly predictive.

Figure S6. Bioinformatic classification allows for accurate OGT prediction.

Figure S7. Genome size is not necessary for OGT prediction accuracy.

Figure S8. Excluding psychrophiles improves OGT prediction.

Figure S9. OGT prediction validated using previously unknown species-OGT values.

Equation S1. Features and coefficients for the prediction of the OGT for a prokaryote.

Equation S2. Features and coefficients for the prediction of the OGT for an Archaea.

Equation S3. Features and coefficients for the prediction of the OGT for a Bacterium.

Table S1. Correlation of features to OGT.

Table S2. De *novo* predicted OGT for species without a measured OGT in Sauer *et al.* 2015

